# An endothelial specific mouse model for the capillary malformation mutation Gnaq p.R183Q

**DOI:** 10.1101/2025.03.17.643802

**Authors:** Patrick J. Smits, Leanna Marrs, Yu Sheng Cheng, Michal Ad, Sana Nasim, David Zurakowski, Joyce J. Bischoff, Arin K. Greene

**Affiliations:** Department of Plastic & Oral Surgery, Boston Children’s Hospital, Harvard Medical School, Boston, MA, 02115, USA; Department of Anesthesiology, Critical Care and Pain Medicine Research, Boston Children’s Hospital and Harvard Medical School, MA. 02115, USA; Vascular Biology Program, Boston Children’s Hospital, Boston, MA, 02115, USA

**Keywords:** capillary malformation, Cdh5Cre, Cdh5CreER, conditional R26 allele, endothelial, GNAQ pR183Q, murine, disease model

## Abstract

Capillary malformation (CM) is a congenital, non-hereditary lesion composed of enlarged and tortuous blood vessels. CM is associated with a somatic GNAQ p.R183Q activating mutation in endothelial cells (EC). Cutaneous CMs are present in 1/300 infants and in 55-70% of CM cases soft tissue overgrowth is observed. Pharmacotherapy for CM does not exist. Here we report a conditional mouse model allowing the simultaneous tissue specific expression of GNAQ p.R183Q and GFP from the *R26* locus (*R26^GT-Gnaq-GFP^*). We show that expression of GNAQ p.R183Q in ECs results in vascular malformations with features similar to human CM lesions. GNAQ p.R183Q expression during embryonic development (*Tg-Cdh5Cre)* resulted in a severe vascular phenotype, lethal by embryonic (E) 16.5. Induction of mutant GNAQ expression in ECs at postnatal (P) day 1 (*Tg-Cdh5CreER)* led to tortuous and enlarged blood vessels, most noticeable in the intestines. GNAQ p.R183Q/GFP expressing ECs co-localized with lesions and displayed increased proliferation. Mutant ECs had abnormal mural cell coverage and abnormal pericellular extracellular matrix deposition, which was confirmed in human CM samples. Similar to human CM they displayed strong expression of the tip cell marker ESM1 and increased ANGPT2 expression. In conclusion, GNAQ p.R183Q expression in murine ECs causes vascular malformations supporting the causality of the mutation for CM. The lesions recapitulate multiple features of human CM, making the mouse model suitable for the preclinical testing of future CM pharmacotherapy.

## INTRODUCTION

Capillary malformation (CM) (also called “port-wine birthmark”) is a congenital sporadic vascular anomaly. It is the most common type of vascular malformation, affecting 1/300 persons[1]. CMs present at birth as pink cutaneous birthmarks that can be located on any area of the body. CM lesions are characterized histologically by widened capillaries with irregular shapes. The dilated vessels have increased pericyte coverage, a thickened basement membrane, and disorganized collagen and elastic fibers, resulting in a thicker blood vessel wall[2, 3]. CMs darken over time and can cause overgrowth of the tissues beneath the birthmark[4]. The deformity caused by CM can lead to functional impairment depending on its location and psychosocial morbidity from the appearance of the lesion. A facial CM in the V1/V2 trigeminal nerve distribution is often associated with Sturge-Weber syndrome (SWS), which also includes ocular and/or brain vascular malformations. The abnormal brain and choroid vasculature in SWS can cause seizures, cortical atrophy, varying levels of cognitive function, glaucoma, and vision loss[5, 6]. Currently, pharmacotherapy for CM/SWS does not exist and primary treatment for CM consists of pulsed-dye laser to lighten the color of the lesion and surgical resection of overgrown tissues. However, CMs often re-darken and re-grow after these interventions[7, 8].

A majority of CM/SWS patients carry somatic single nucleotide changes in *GNAQ*, with c.548G>A; p.R183Q being the most common mutation. A subset of CM/SWS patients carry a single nucleotide change at the same conserved site, p.Q183 in the closely related GNA11 gene[9–12]. The GNAQ p.R183Q change is enriched in the endothelial cells (ECs) of skin, brain and eye lesions in both CM and SWS, but has also been found at low frequency in non-ECs[10, 13, 14]. GNAQ, together with GNA11,14 and 15, is a member of the Gq-alpha protein family[8]. It encodes Gαq, the alpha subunit of the heterotrimeric Gq protein which relays signals from transmembrane G-protein coupled receptors (GPCRs) to phospholipase Cβ (PLCβ). Several vascular GPCRs are known to signal through Gq including protease-activated-receptor-1 (PAR1)[15], bradykinin receptors[15], endothelin receptors[16] and angiotensin-II type I receptor[17].

Inactive, GDP bound, Gαq is anchored to the cell membrane in a protein complex with the β and γ subunits. Upon GPCR ligand binding, Gαq exchanges GDP for GTP. This leads to dissociation of the heterotrimeric protein complex, with the GTP-bound active Gαq able to initiate downstream signaling. The Gαq subunit, which has intrinsic GTPase activity, remains active until its GTP is hydrolyzed back to GDP allowing the reformation of the heterotrimeric protein complex. The CM associated point mutations in GNAQ are located in its so-called switch I region, alter its structure and lower Gαq’s GTPase activity. This leads to a higher percentage of the protein being in its active GTP bound form, increasing downstream signaling through phosphorylation of PLCβ. Activated PLCβ catalyzes the production of inositol trisphosphate (IP_3_) and diacylglycerol (DAG). IP_3_ triggers cytoplasmic calcium release which together with DAG activates PKC signaling. PKC on its turn can both activate MAPK/ERK signaling and promote NF-κB nuclear translocation. The NF-κB activation has been shown to increase the expression of ANGPT2, a potent angiogenic factor[8, 9, 18–21].

Our ability to understand how CM and SWS form and grow over time and the development of effective pharmacotherapy has been hampered by the absence of suitable mouse models. Specific expression of GNAQ p.Q209L, the activating mutation associated with uveal melanoma and a subset of vascular tumors, but not CM or SWS, in postnatal murine ECs was shown to cause vascular abnormalities[22]. A conditional murine model allowing expression of GNAQ p.R183Q from the endogenous *Gnaq* locus has also been reported[23]. However, this study did not use EC specific expression of the mutant protein but instead induced broad mosaic GNAQ p.R183Q expression. No postnatal vascular phenotype was present, but dilated blood vessels were observed in 20% of mutant embryos. More recently, a Tet-ON GNAQ p.R183Q transgenic mouse line was reported[24]. When bred to Tie2-rtTA tet trans activator mice this line allows EC specific CMV promotor driven overexpression of GNAQ p.R183Q upon exposure of the mice to doxycycline. This study focused on SWS and the leptomeningeal vasculature of the mutant mice. Blood-brain barrier breakdown and abnormal cortical micro vessel structure was reported.

Here we report the generation of a conditional *Rosa26 Gnaq* p.R183Q murine model allowing simultaneous expression of GNAQ p.R183Q and GFP from the same transcript. We show, using Tg-*Cdh5Cre* and Tg-*Cdh5CreER*, that EC specific expression of GNAQ p.R183Q generates vascular malformations embryonically and postnatally in all tissues. Post-natal, we mimicked the somatic nature of the CM mutation by activating GNAQ p.R183Q expression in only a subset of ECs. We present a detailed characterization of the murine lesions showing that they recapitulate multiple features of human CM. This animal CM model will further enable study into how GNAQ p.R183Q changes EC behavior and drives CM formation and progression. It also will be a useful preclinical model for the development of CM targeted medications.

## RESULTS

### Generation of a conditional R26^GT-Gnaq-GFP^ allele

To generate a mouse model allowing for the conditional expression of GNAQ p.R183Q in ECs we used CRISPR/Cas9 gene editing to introduce a p.R183Q mutant *Gnaq* cDNA into the *R26* locus. The targeting vector contained a splice acceptor (SA), preventing expression from the endogenous *R26* promoter, upstream of a strong synthetic *CAG* promoter. The *CAG* promoter was followed by a LoxP flanked stop cassette (gene trap (GT)). The *Gnaq p.R183Q* cDNA was cloned 3’ prime of the stop cassette and was followed by an independent ribosomal entry site (IRES)-GFP cassette. After removal of the GT the *CAG* promoter drives expression of a bis-cistronic *Gnaq p.R183Q-GFP* mRNA leading to expression of both GNAQ p.R183Q and GFP. Expression of GFP allows us to identify cells containing a recombined allele (**Fig. 1A**).

**Figure 1:**
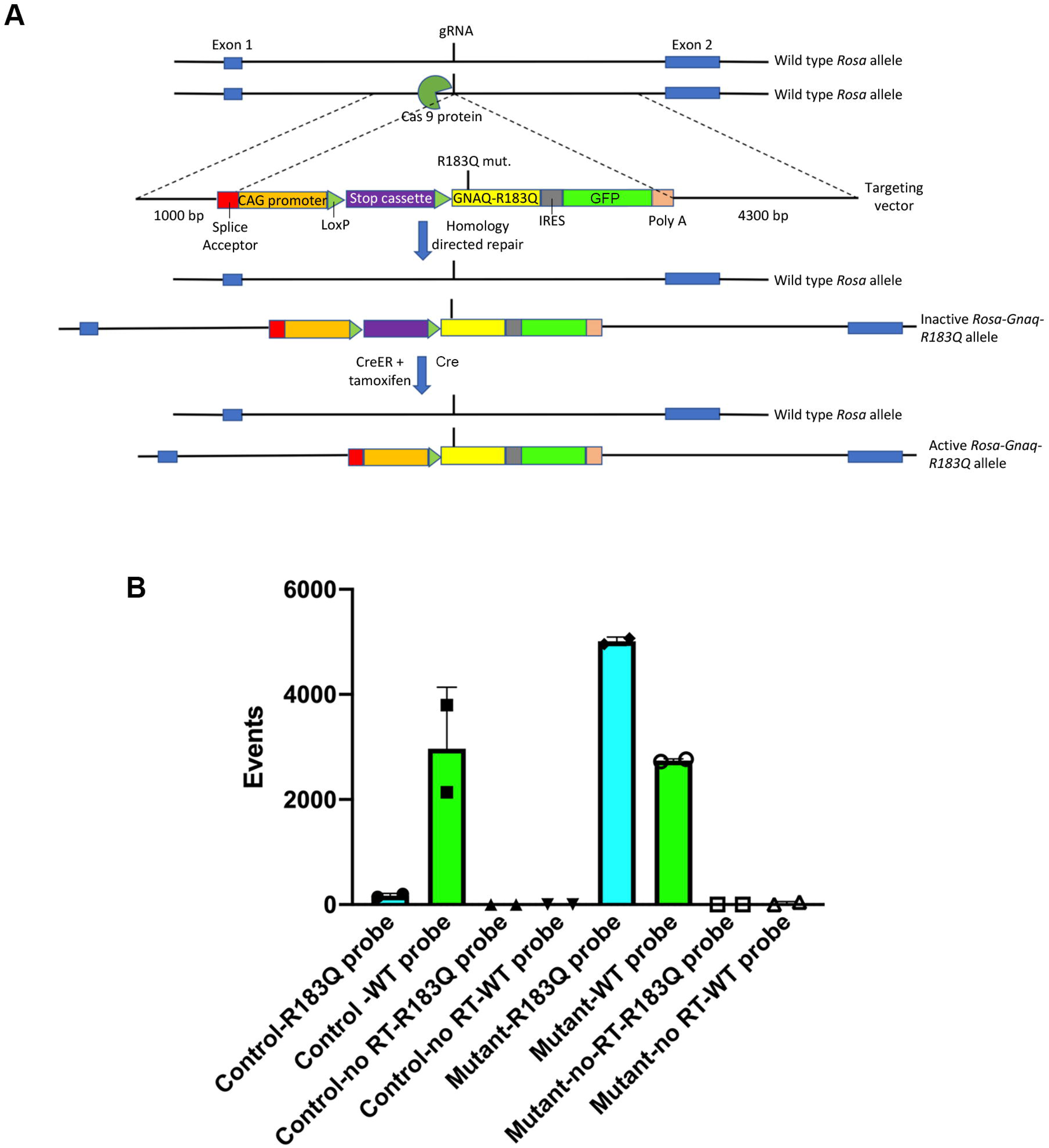
Schematic of the conditional R26^GT-Gnaq-GFP^ allele. **(A)** The LoxP flanked gene trap prevents expression of the allele before recombination. The IRES-GFP sequence allows identification of cells with an active allele. **(B)** High expression of *Gnaq* p.R183Q after gene trap removal. ddPCR using cDNA generated from tamoxifen treated lung ECs. Control: *R26^GT-Gnaq-GFP^,* mutant: *Tg-Cdh5CreERT2^+/-^;R26^GT-Gnaq-GFP/+^*. Probes used: *Gnaq* p.R183Q (blue) and *Gnaq* wild type (green).

To confirm the functionality of the *R26^GT-Gnaq-GFP^* allele after recombination, we isolated lung ECs from 6-week-old *Tg-Cdh5CreERT2^+/-^;R26^GT-Gnaq-GFP/+^* and *R26^GT-Gnaq-GFP/+^* mice, cultured them in the presence of 4-OH-tamoxifen, and extracted RNA. ddPCR with WT *Gnaq* and *Gnaq p.R183Q* specific probes using cDNA, generated from the lung EC RNA, showed that 68% of *Gnaq* transcripts in tamoxifen treated *Tg-Cdh5CreERT2^+/-^;R26^GT-Gnaq-GFP/+^* lung ECs had the R183Q mutation (**Fig. 1B**). Low expression of *Gnaq p.R183Q* (5.75% of *Gnaq* transcripts) was detected in *R26^GT-Gnaq-GFP/+^* lung ECs indicating that the gene trapped allele has low level leakage. However, this leakage did not produce vascular or other anomalies, and *R26^GT-Gnaq-GFP/+^* animals were healthy, fertile, and had normal life spans.

### Expression of GNAQ p.R183Q during embryonic development is lethal

To obtain activation of the *R26^GT-Gnaq-GFP/+^* allele in ECs during embryonic development we used the *Tg-Cdh5Cre* transgene. This transgene has been reported to drive pan EC expression by embryonic day (E) 14.5[25]. Genotyping of neonatal offspring from breeding a *Tg-Cdh5Cre^+/+^* sire (homozygous for the Cre transgene) with *R26^GT-Gnaq-GFP/+^* females showed the absence of *Tg-Cdh5Cre^+/-^;R26^GT-Gnaq-GFP/+^* pups (0 out of 33 pups genotyped), indicating that pan EC expression of GNAQ p.R183Q is embryonic lethal. To determine at what time point lethality occurs, we extracted embryos at E13.5, E14.5, E15.5 and E16.5. *Tg-Cdh5Cre^+/-^;R26^GT-Gnaq-GFP/+^*embryos were found at all 4 stages. At E13.5 they were indistinguishable from control littermates (N=5). By E14.5, the vascular network in the developing skin of mutant fetuses (N=9) was visibly different from controls, displaying abnormal vessels on the torso **(Fig. 2A)**. This phenotype worsened by E15.5 (**Fig. 2B**) (N=7). No viable mutant embryos were recovered at E16.5 (N=5). Histological analysis of E14.5 embryos showed the presence of enlarged blood vessels which were most prominent in the developing skin **(Fig. 2C).** Hemorrhage occasionally occurred, most notable into the jugular lymph sacks. (**Fig. 2D**).

**Figure 2:**
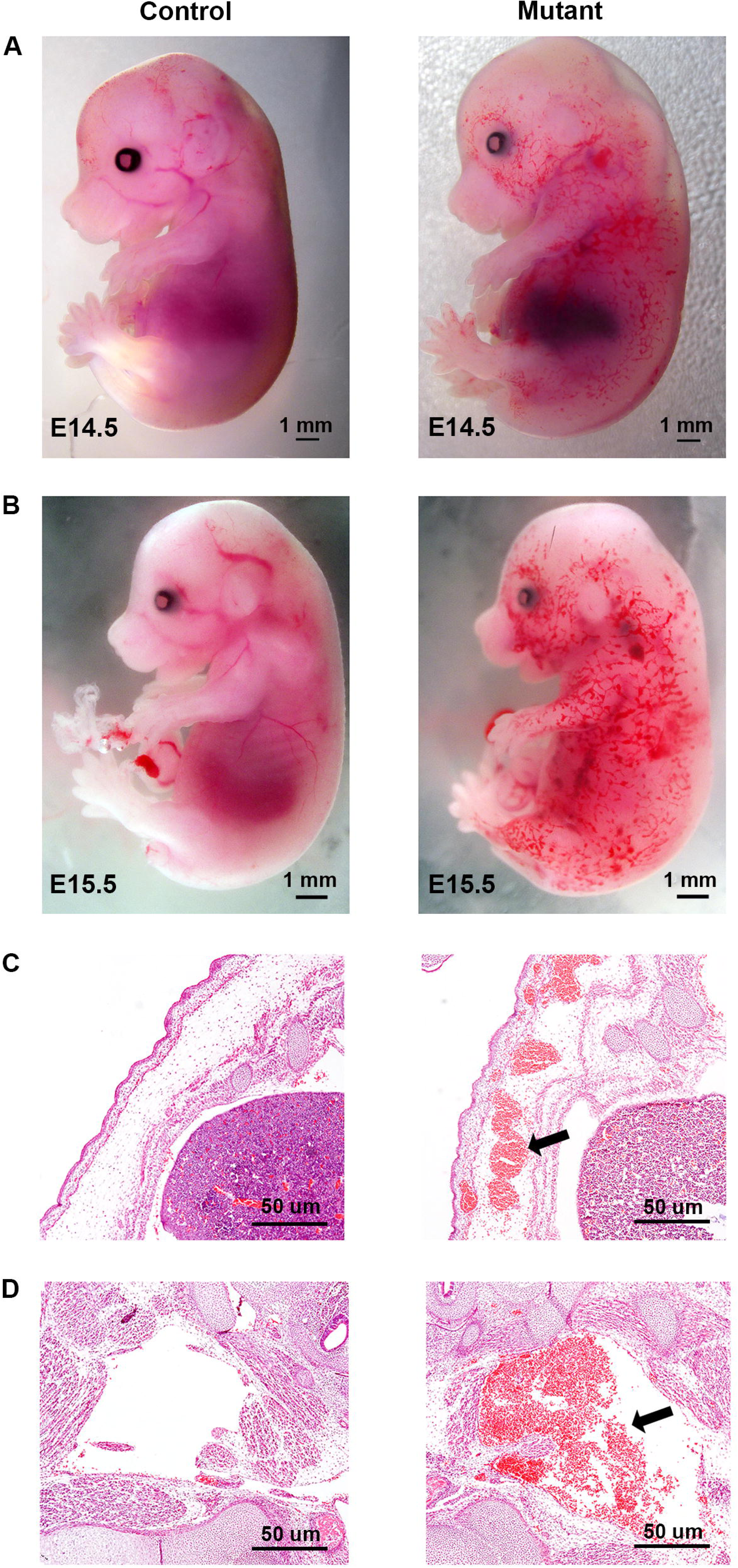
In utero EC GNAQ p.R183Q expression is lethal. **(A)** Phenotype of E14.5 *Tg- Cdh5Cre^+/-^*(control) and *Tg-Cdh5Cre;R26^GT-Gnaq-GFP/+^* (mutant) embryos. Note diffuse abnormal vasculature on the torso of the mutant. **(B)** Phenotype of E15.5 embryos. The mutant embryo displays more severe vascular abnormalities. (**C)** H&E-stained developing ventral skin of E14.5 embryos. shows enlarged blood vessels (arrows) in the mutant embryo. **(D)** H&E-stained developing left jugular lymph sacks of E14.5 embryos. The jugular lymph sack in the mutant embryo is filled with blood (arrow).

#### Neonatal activation of Gnaq p.R183Q expression in ECs results in vascular abnormalities

To circumvent the embryonic lethality, we used the *Tg-Cdh5CreERT2* transgene. *Tg- Cdh5CreERT2^+/-^;R26^GT-Gnaq-GFP/+^* and control littermates were subcutaneously injected with tamoxifen at P1 **(Fig. 3A)**. Injection of 100 µg of tamoxifen resulted in growth retardation and lethality between P15-P37 (N=14) **(Fig. 3B)**. Animals did not deteriorate over time but died suddenly. Necroscopy showed widespread vascular anomalies most obvious in skeletal muscles and the gastrointestinal tract. Histological analysis of the intestinal submucosa showed that the anomalies contained enlarged blood vessels **(Fig. 3C)**. No visible phenotype was present in the brain, however GFP expression could be detected on the brain surface of *Tg-Cdh5CreERT2^+/-^;R26^GT-Gnaq-GFP/+^*animals (**Fig. 3D**). Because GFP is translated from the second open reading frame of the single *R26^GT-Gnaq-GFP^* transcript, green fluorescence marks ECs with an activated *R26^GT-Gnaq-GFP/+^* allele. Sectioning through the brain showed the presence of enlarged blood vessels in the leptomeninges of mutant animals (**Fig. 3D**).

**Figure 3:**
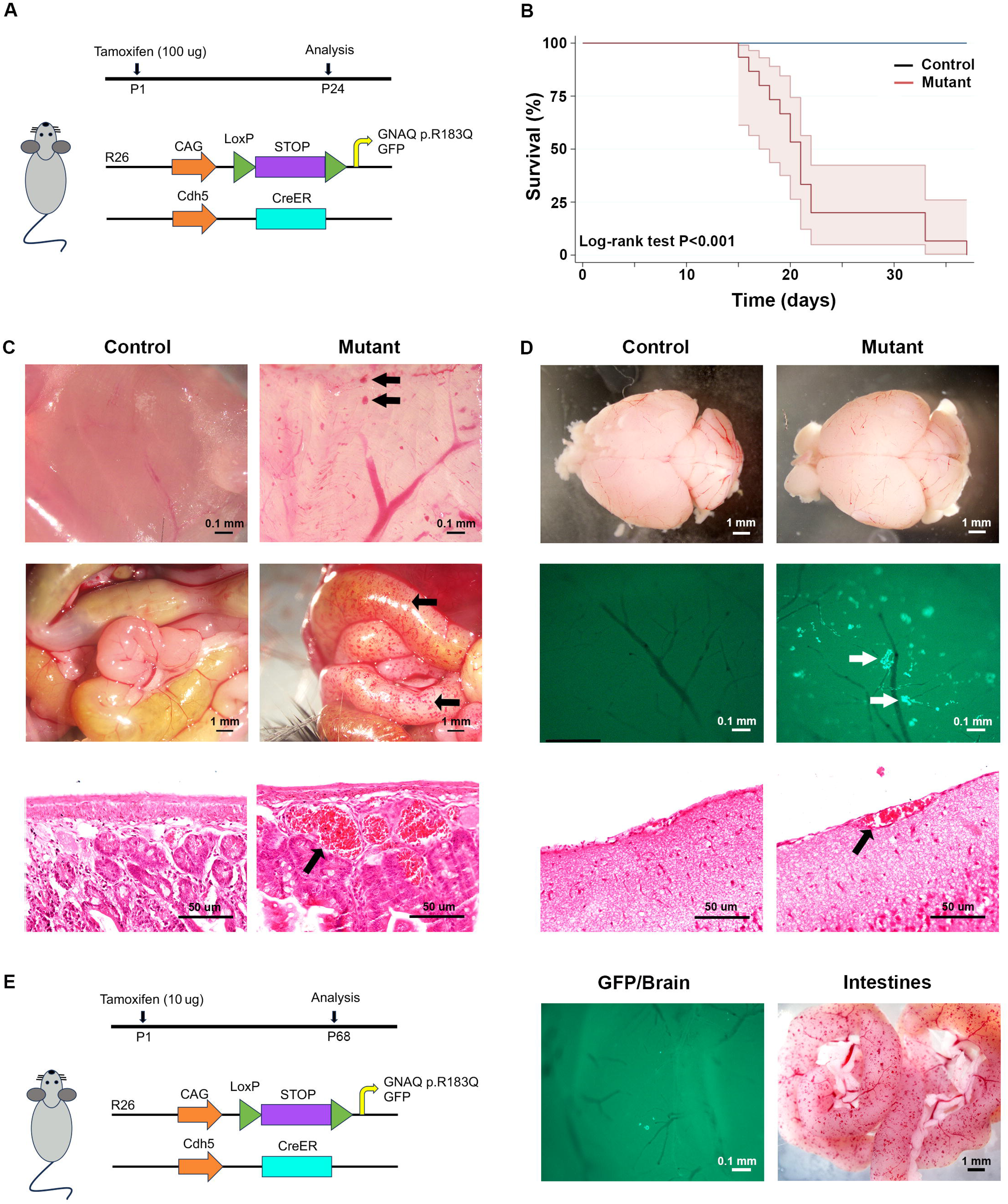
Neonatal expression of Gnaq p.R183Q in ECs results in vascular malformations. **(A)** Tamoxifen induction scheme. *R26^GT-Gnaq-GFP/+^* females were mated with *Tg-Cdh5CreER^+/-^ sires.* Offspring was treated with 100 µg of tamoxifen at P1. Phenotypic analysis was done at P24. **B)** Kapler Meier survival curves of tamoxifen treated *Tg-Cdh5CreER^+/-^;R26^GT-Gnaq-GFP/+^* (N=15) and control animals *(WT, Tg- Cdh5CreER^+/-^ or R26^GT-Gnaq-GFP/+^)* (N=14). **(C)** Gross phenotype of P24 mice treated with tamoxifen at P1. Left, *Tg-Cdh5CreER^+/^*^-^ (control). Right *Tg-Cdh5CreER^+/-^;R26^GT-Gnaq-GFP/+^*(mutant). Top panels: pectoral muscle. Middle panels: intestines. Bottom panels: H&E- stained sections through the intestines. Note the abnormal vasculature in the mutant muscle and intestines (arrows). (**D**) Brains of P1 tamoxifen treated P24 mice. Top panels: gross appearance. Middle panels: fluorescence imaging of GFP. Note the presence of GFP positive cells on the surface of the brain (arrows). Bottom panels: H&E-stained sections. Note enlarged blood vessels in the mutant leptomeninges (arrows). **(E)** Phenotype of aged mice treated with a lower dose of tamoxifen. Left panel: Tamoxifen induction scheme. Mice were injected at P1 with 10 µg of tamoxifen (N=11). Phenotypic analysis was done at >2 months. Right panels: GFP detection in brain and gross intestinal phenotype. Images are from a P68 mouse. Note the fewer GFP expressing vessels in the brain but a similar severity of the intestinal phenotype.

Injection of P1 *Tg-Cdh5CreERT2^+/-^;R26^GT-Gnaq-GFP/+^* pups with 5 µg of tamoxifen increased the life span of the animals to >2-months (N=11). These animals did not display growth retardation or sudden death. Necroscopy revealed fewer GFP expressing ECs in the brain of the animals while the gastrointestinal phenotype was unaffected by the lower tamoxifen dose **(Fig. 3E)**.

Whole mount immunofluorescence staining of the intestines and dorsal ear skin of P1 tamoxifen treated *Tg-Cdh5CreERT2^+/-^;R26^GT-Gnaq-GFP/+^*animals for CD31 and GFP confirmed the sporadic activation of the *R26^GT-Gnaq-GFP/+^*allele, with only a percentage of ECs expressing GFP. Furthermore, vessel abnormalities coincided with the presence of GNAQ p.R183Q expressing (GFP positive) ECs **(Fig. 4A-B)**. Mutant tissues displayed two types of vessel abnormalities, vessel bulging and abnormal vessel tortuosity **(Fig. 4C)**.

**Figure 4:**
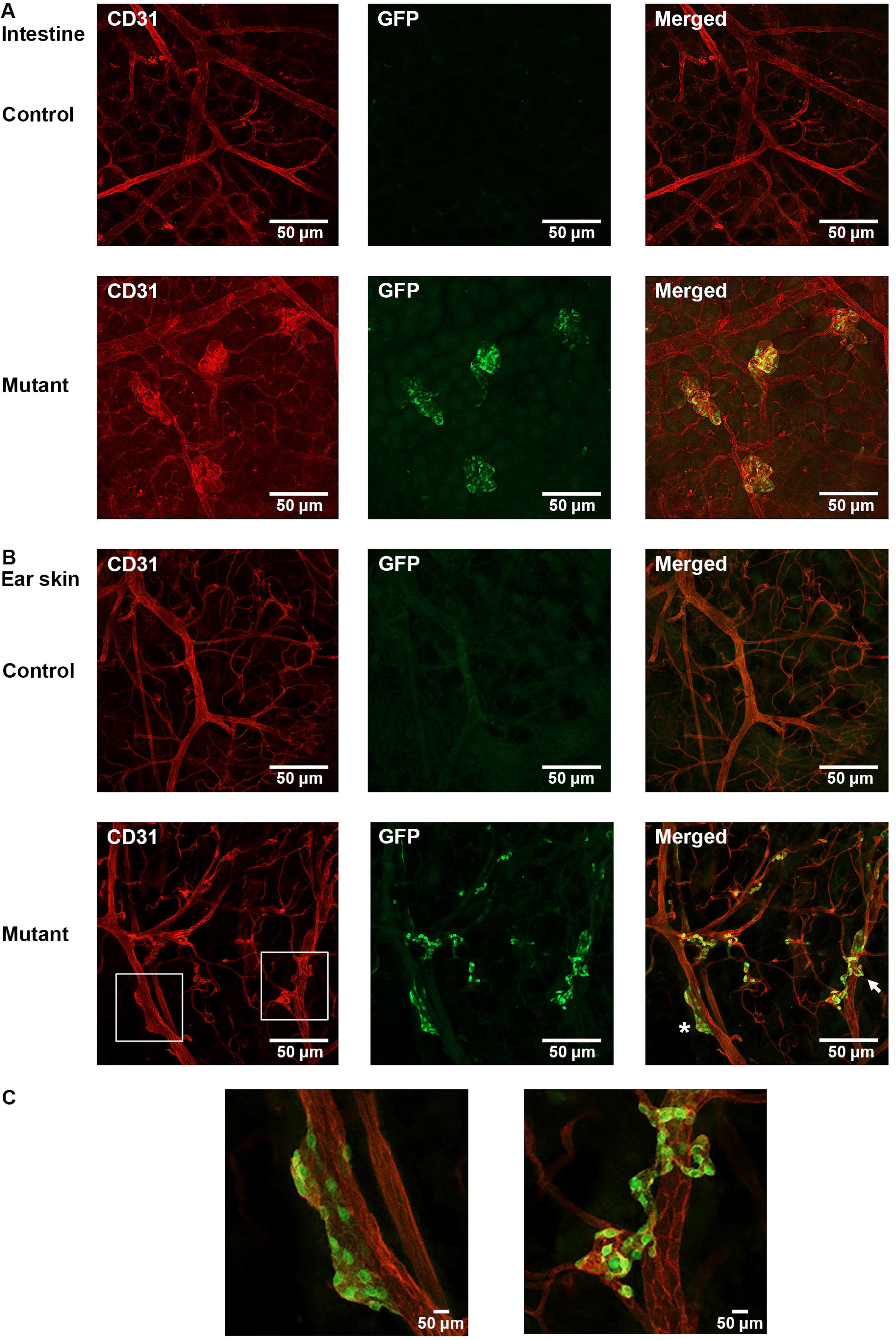
Gnaq p.R183Q expressing cells co-localize with vessel malformations. Single channel and merged images of immunofluorescent staining with CD31 (red) and GFP (green) of the intestines **(A)** and dorsal ear skin **(B)** from P24 mice treated with tamoxifen at P1. Top panels: *Tg-Cdh5CreER^+/^*^-^ (control). Bottom panels*: Tg-Cdh5CreER^+/-^;R26^GT-Gnaq-GFP/+^* (mutant). Note that GFP expressing cells co-localize with areas of blood vessel enlargement (asterisk) and tortuosity (arrow). **(C)** Higher magnification of the enlarged and tortuous vessels marked in B.

#### GNAQ p.R183Q expression increases EC proliferation and induces an EC tip cell identity

Increased EC proliferation due to hyperactive GNAQ signaling and increased sprouting have been associated with the development of CMs. Activation of GNAQ signaling leads to increased melanocyte proliferation in uveal melanoma (caused by the activating GNAQ p.Q209L mutation)[26], while ECs in enlarged vessels of human CM have been shown to express the tip cell marker ESM1[27]. To determine whether these features are also present in our mouse model after induction of GNAQ p.R183Q expression, we performed whole mount immuno-fluorescence staining of the intestines of tamoxifen treated *Tg-Cdh5CreERT2^+/-^;R26^GT-Gnaq-GFP/+^*animals (N=3) for the proliferation marker Ki67, the EC specific transcription factor ERG (to identify ECs), GFP (to identify GNAQ p.R183Q expressing ECs) and ESM1. ECs were found to have a low proliferation rate with few ERG+ cells staining for Ki67. However, a significant difference in the number of Ki67+ cells was observed between mutant ECs (ERG+/GFP+: 3.5% average) and non-mutant ECs (ERG+/GFP-: 0.61% average) **(Fig 5A)**. Analysis of EC proliferation in tamoxifen treated control animals (*R26^GT-Gnaq-GFP/+^*) (N=2) showed a similar proliferation rate as the ERG+/GFP- EC population of mutant animals. ERG staining further showed that mutant EC nuclei within the lesions are round compared to the flattened elongated nuclei shape of WT ECs. Analysis of ESM1 expression in mutant animals showed overlap with GFP positive lesions, indicating that GNAQ p.R183Q expressing ECs have a tip cell identity **(Fig. 5B)**.

**Figure 5:**
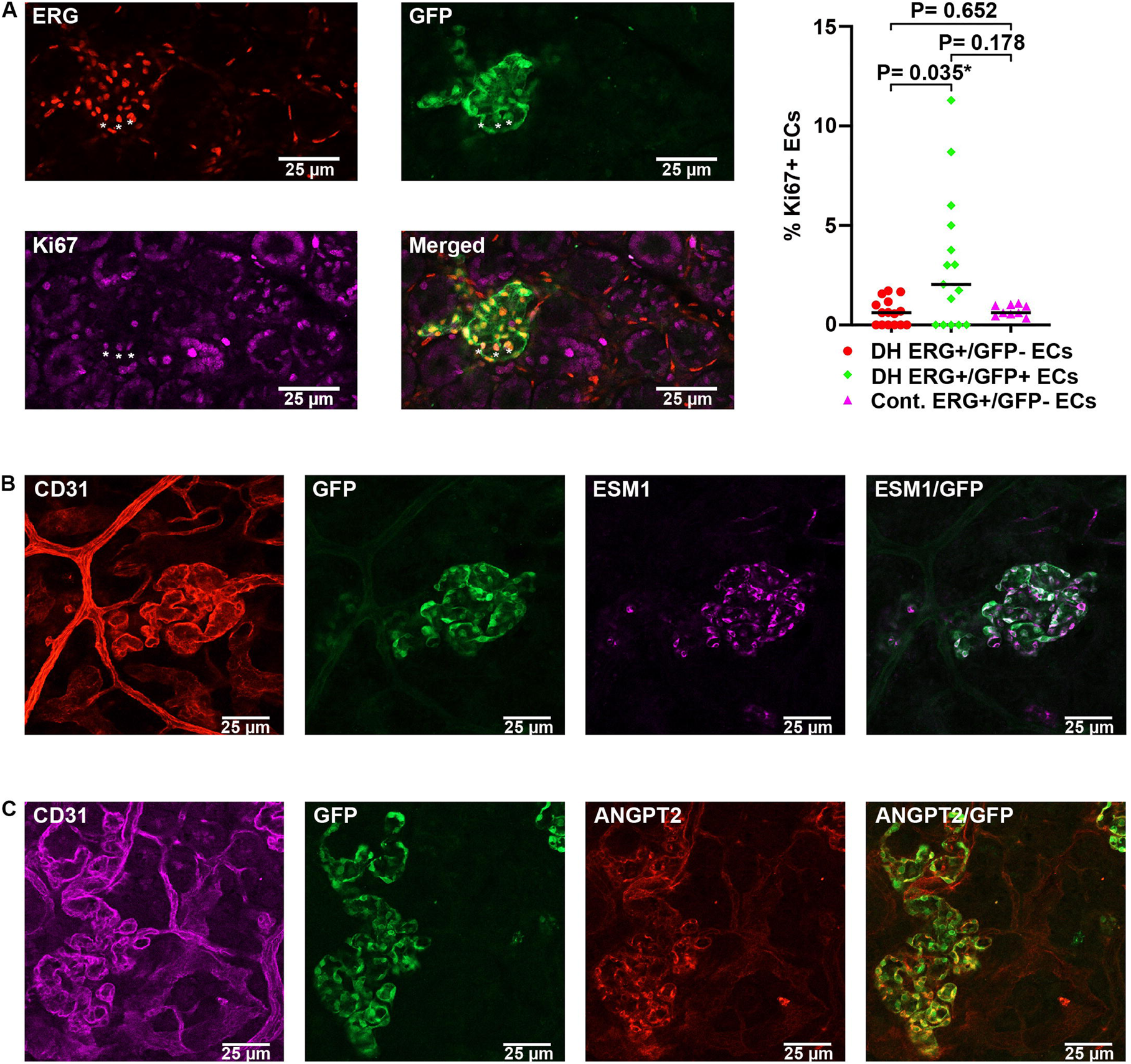
Gnaq p.R183Q expression increases EC proliferation and induces a tip cell identity. **(A)** Left panels: representative single channel and merged images of immunofluorescent staining of mutant intestines for CD31, Ki67 and GFP. * Marks triple positive ECs within the lesion. Right panel: graph comparing the percentage of Ki67 positive ECs within the lesions (ERG+/GFP+) with non-mutant ECs outside the lesions (ERG+/GFP-) and ECs of control (*R26^GT-Gnaq-GFP/+^*) animals. GNAQ p.R183Q expressing ECs have a higher proliferation rate. **(B)** Single channel and merged images of immunofluorescent staining of mutant intestines for CD31, GFP and ESM1. Note that ESM1 all GFP+ ECs stain for ESM. **(C)** Single channel and merged images of immunofluorescent staining of mutant intestines for CD31, GFP and ANGPT2. Note that staining for ANGPT2 staining is stronger in the GFP positive lesion.

### Increased ANGPT2 expression in GNAQ p.R183Q ECs

An increase in the levels of the angiogenic factor ANGPT2 in GNAQ p.R183Q expressing HUVECs has previously been reported[21]. To confirm whether this is recapitulated in our mouse model we performed whole mount immunofluorescence staining of the intestines of *Tg-Cdh5CreERT2^+/-^;R26^GT-Gnaq-GFP/+^*animals, treated with tamoxifen at P1, for ANGPT2. GFP positive vascular lesions showed stronger ANGPT2 staining compared to non-GFP expressing blood vessels **(Fig. 5C)**.

### Gnaq p.R183Q expressing ECs have increased pericellular extracellular matrix deposition

The pericellular matrix, which is produced by both ECs and mural cells, has multifaceted roles in EC biology. It is a dynamic environment providing not just a scaffold for EC anchoring but is also involved in regulating blood vessel diameter and integrity, EC migration, and the availability of growth factors[28]. Actively sprouting ECs change their pericellular matrix[29]. Human CM is characterized by a thickening of the basement membrane of affected blood vessels[3]. To study whether pericellular matrix deposition of GNAQ p.R183Q ECs is affected in our mouse model, we co-stained the intestines of tamoxifen treated *Tg-Cdh5CreERT2^+/-^;R26^GT-Gnaq-GFP/+^* animals for CD31, GFP and pericellular matrix components. We found that staining for the basement membrane proteins HSPG2, NIDOGEN2, and COLLAGEN IV was increased around GFP positive ECs **(Fig. 6A)**. Staining of human CM tissue sections confirmed these findings, with abnormally enlarged blood vessels surrounded by a thicker layer of COL IV expressing cells **(Fig 6B).**

**Figure 6:**
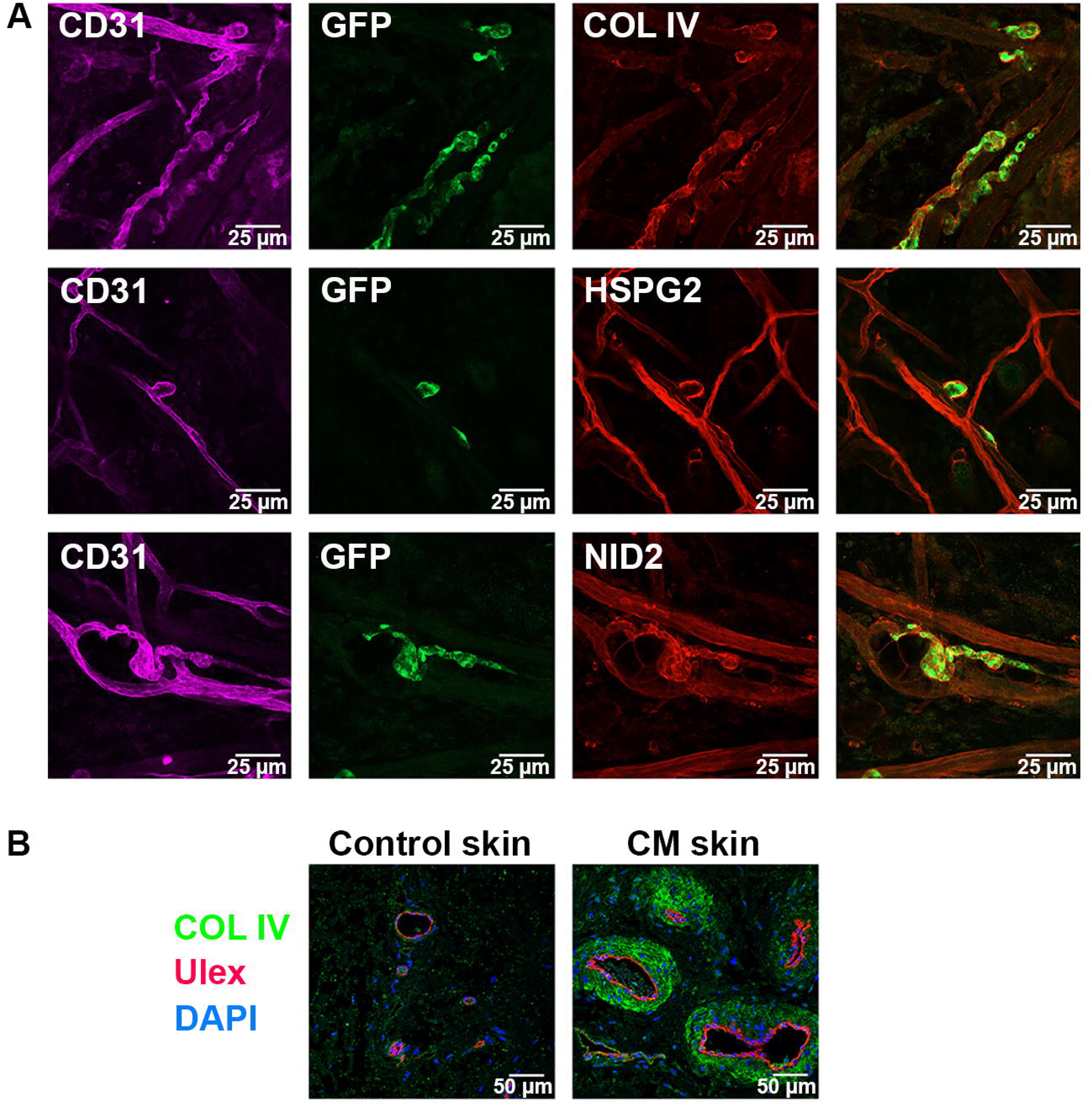
GNAQ p.R183Q lesions display increased pericellular matrix deposition. **(A)** Single channel and merged images of immunofluorescent staining of mutant intestines for CD31, GFP and the basement membrane components COLLAGEN IV, HSPG2 and NIDOGEN-2. Note the stronger staining for the basement membrane proteins around GFP positive ECs. **(B)** Immunofluorescent staining of human CM and control skin for COLLAGEN IV illustrates broader staining (green) around enlarged vessels.

### GNAQ p.R183Q expression impairs EC-mural cell interaction

Mural cells – pericytes and vascular smooth muscle cells – are integral to maintaining functional blood vessels. They provide structural support, regulate blood flow (modulating vascular diameter and contractility), reduce vessel leakage, contribute to the blood vessel extracellular matrix and play roles in vessel maturation and patterning[30]. A feature of human CM is disorganized mural cell coverage of enlarged vessels[3, 27]. We therefore analyzed whether inducing EC expression of GNAQ p.R183Q in our mouse model results in similar changes. Whole mount immunofluorescence staining of intestines of tamoxifen treated *Tg-Cdh5CreERT2^+/-^;R26^GT-Gnaq-GFP/+^* animals for CD31, GFP and the pericyte markers NG2 and PDGFRb showed that GFP positive lesions featured more intense NG2 and PDGFRb staining **(Fig. 7A)**. Staining for the vascular smooth muscle cell marker alpha-SMA showed that GFP positive regions of blood vessels occasionally contained both alpha-SMA positive and alpha SMA negative ECs. Immunofluorescent staining of human CM tissue sections showed a broader layer of NG2 positive pericytes surrounding enlarged blood vessels, confirming the mouse data **(Fig. 7B).**

**Figure 7:**
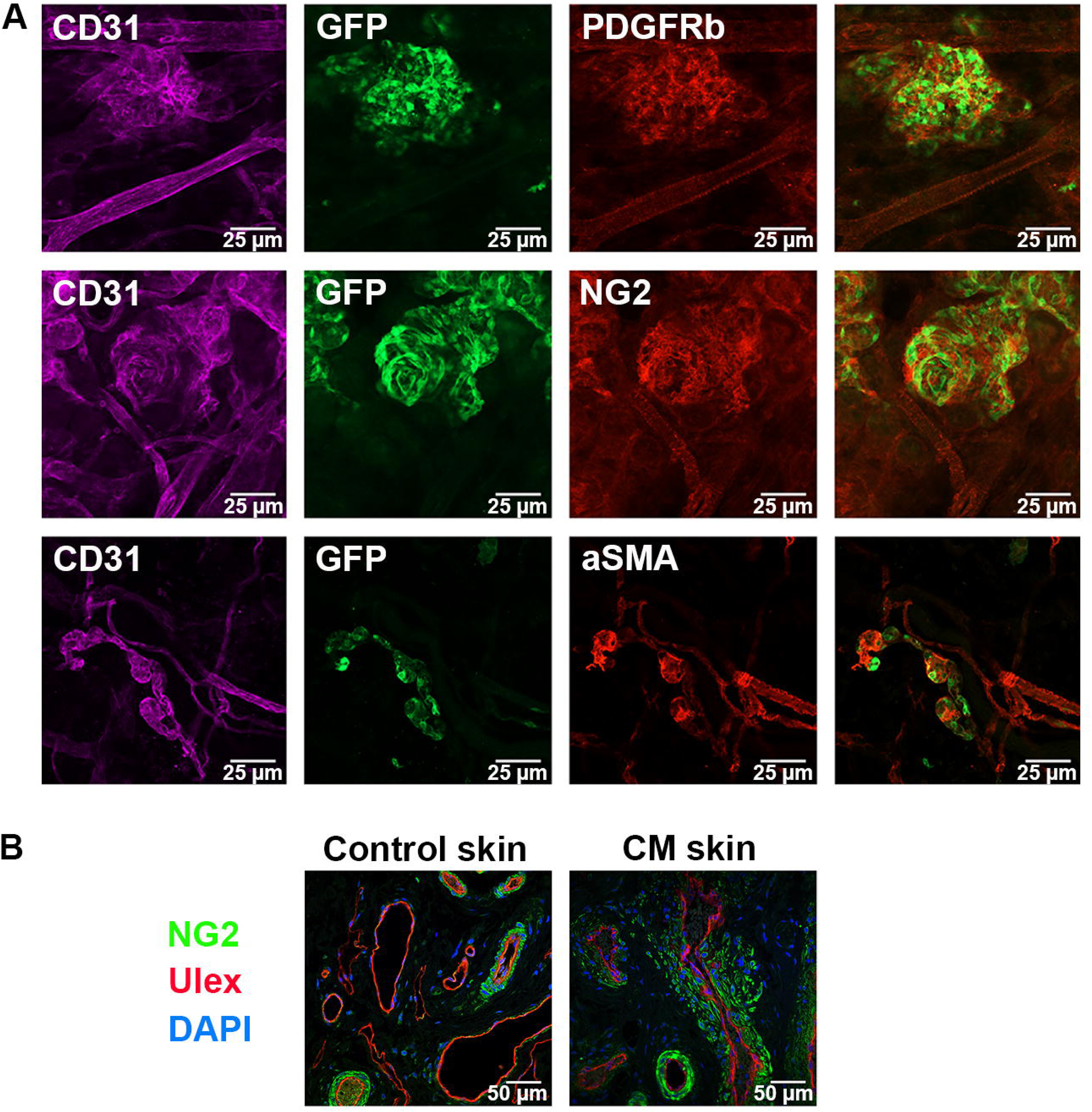
GNAQ p.R183Q expressing ECs display impaired mural cell interactions. **(A)** Single channel and merged images of immunofluorescent staining of mutant intestines or ear skin for CD31, GFP and the mural cell markers PDGFRb, NG-2 and alpha-SMA. Note that intestinal lesions stain stronger for PDGFRb and NG-2, while alpha-SMA staining is irregular in the GFP positive dorsal ear skin vessel. **(B)** Immunofluorescent staining of human CM and control skin for NG-2. Note that enlarged vessels have multiple layers of NG-2 expressing pericytes.

## DISCUSSION

A somatic activating *Gnaq* p.R183Q mutation is the most common cause of CM/SWS[9, 11]. We developed a murine model allowing for the conditional expression of GNAQ p.R183Q from the R26 locus. Using the endothelial specific *Tg-Cdh5Cre* and *Tg- Cdh5CreER* transgenes we generated mice expressing mutant GNAQ protein in ECs. We found that pan-EC GNAQ p.R183Q expression during embryonic development is incompatible with life, illustrating the importance of GNAQ signaling for the development and maintenance of the vascular system. By exposing neonatal *Tg-Cdh5CreERT2^+/-^;R26^GT-Gnaq-GFP/+^* animals to low doses of tamoxifen, we were able to recapitulate the human CM paradigm that the CM/SWS mutation is somatic and enriched in ECs. Somatic GNAQ p.R183Q EC expression resulted in diffuse vascular malformations likely in all vascularized tissues. The vascular lesions are most prominent in the intestines with affected vessels featuring bulging and increased tortuosity. Designing the conditional allele to express both GNAQ p.R183Q and GFP from the same transcript allowed us to identify ECs expressing GNAQ p.R183Q based on green fluorescence. We found that mutant ECs localized to affected regions and that lesions were made up exclusively of GFP expressing ECs. These results confirm that mutant ECs are the driving force behind CM/SWS lesion formation. They also show that the *Gnaq* p.R183Q mutation in ECs is sufficient to cause vascular malformations in mice, supporting *GNAQ* p.R183Q pathogenesis in human CMs.

To validate our mouse model, we cross checked its phenotype with known features of human CM. Histological analysis of the mutant intestines and brain showed blood vessels with an enlarged diameter, similar to what is observed in human CM/SWS[8, 9]. Furthermore, we found that GNAQ p.R183Q impairs normal EC-mural cell interaction and affects the EC pericellular matrix. Based on marker expression, lesions displayed increased pericyte coverage and basement membrane deposition. Both findings have been associated with CM[3], and we confirmed them in patient CM skin samples. Other human CM features recapitulated in our mouse model are increased EC proliferation, increased ANGPT2 levels and, the induction of expression of the tip cell marker ESM1 in GNAQ p.R183Q ECs[9, 21, 27]. We also observed that the nuclei of mutant ECs in intestinal lesions have a rounded appearance. ECs normally align with the laminar blood flow and have an elongated appearance with flatter more stretched out nuclei. The rounded nuclei in GNAQ p.R183Q ECs suggest that these cells do not respond correctly to laminar flow, a feature that has also been reported for human GNAQ p.R183Q EC models[31].

Unlike human CMs, which mainly affect the integument, our mice did not show any grossly visible skin lesions. Despite the fact that mutant ECs are present and microscopic vascular malformations can be observed in the skin, our *Tg-Cdh5CreERT2^+/-^;R26^GT-Gnaq-GFP/+^* animals do not develop an extensive vascular network reminiscent of CM, even when exposed to the lower tamoxifen doses that allows them to age (the oldest animals analyzed were >3-months). Furthermore, although GNAQ p.R183Q ECs express the tip cell marker ESM1, we did not observe ECs sprouting out from the lesions. The absence of sprouting could explain why GNAQ p.R183Q ECs despite their increased proliferation rate fail to form a network. Species difference in skin structure could account for the absence of visual skin lesions in mice. For example, mouse skin lacks sweat glands and contains abundant hair follicles. Intriguing is that the GNAQ p.R183Q mutation also has been detected in low frequency in the non-endothelial cell population of CMs[13, 32]. It is therefore possible that in human CM there is a contribution from GNAQ mutant non-ECs that is not present in our EC specific mouse model. The GNAQ p.R183Q mutation in humans occurs prenatally. If the mutation happens at an early embryonic stage in a hemangioblast or a mesodermal progenitor cell, it could give rise to two distinct somatic mutated cell populations, ECs and a mural cell type. More complex mouse models using multiple CreER drivers allowing activation of GNAQ p.R183Q in both ECs and mural cells (e.g. PDGFRβ-CreERT2 for pericytes) would be needed to address this. Another possibility is that CM requires two mutational hits, with a second as yet unknown mutation triggering excessive sprouting.

In addition to our mouse model three other conditional *Gnaq* mutant mouse lines have been recently reported, two GNAQ p.R183Q and one GNAQ p.Q209L. Wetzel-Strong et al., activated Gnaq p.R183Q expression from the endogenous *Gnaq* locus but used the ubiquitous expressed *Tg-E2A-Cre* and *Tg-Actin-Cre* transgenes which did not restrict mutant GNAQ expression to ECs[23]. They showed embryonic lethality, hemorrhage, and dilated capillaries but did not present postnatal data. Solomon et al., generated a mouse model allowing doxycycline inducible CMV promoter driven GNAQ p.R183Q expression in ECs[24]. They focused on SWS and the leptomeningeal vasculature but did not report analysis of other tissues vasculature. They showed a breakdown of the blood brain barrier and increased vessel permeability, but did not report the formation of lesions similar to the lesions observed in our model. Finally, Schrenk et al., used a conditional *R26* allele for a different *GNAQ* variant, *GNAQ* p.Q209L, which is found in uveal melanoma and vascular tumors[22, 33]. They did not study embryonic EC specific expression, but showed vascular abnormalities in the subcutis, brain, retina, and intestines with postnatal GNAQ p.Q209L EC expression. The gross appearance of the malformations in this animal model is similar to the lesions in our model and increases in vessel diameter and EC proliferation were also reported. However, despite using the same *Tg-Cdh5CreER* transgene and a 10 times higher tamoxifen dose (100 µg compared to 10 µg in the study by Schrenk and co-workers), our mice survive longer than mice with EC GNAQ p.Q2O9L expression (P15-37 as compared to P6). Both the R183 and Q209 residues in the GNAQ protein are involved in GTP hydrolysis, resulting in switching off GNAQ activity[8, 14, 34]. While the p.Q209L mutation causes a complete loss of GTPase activity and subsequent constitutive activation of GNAQ signaling, the p.R183Q variant leads to slower GTPase activity and a lower degree of GNAQ activation[18, 19, 35]. Galeffi et al., recently suggested that the pQ209L mutation is not tolerated by ECs even when only present mosaically in utero[35]. This difference in activation strength between the two mutations could explain why our animals live longer despite having a more severe phenotype with GNAQ p.Q209L ECs not surviving long enough to develop vascular lesions. This would also explain why while the GNAQ p.R183Q mutation has been found in uveal melanoma, the GNAQ p.Q209L mutation has as of yet never been associated with CM/SWS[36].

In summary, we have shown that somatic expression of GNAQ p.R183Q in murine ECs is sufficient to generate vascular malformations that recapitulate multiple features of human CM. These findings support the causative role of the GNAQ p.R183Q mutation in the formation of human CMs and SWS. Our animal model allows further investigation into the mechanisms by which CMs develop and can serve as a preclinical model for testing pharmaceuticals aimed at treating CM/SWS.

## METHODS

### Generation of a conditional R26^GT-Gnaq-GFP^ mouse line

Animal experiments were approved by Children’s Hospital Boston Institutional Animal Care and Use Committee. All experiments were performed in accordance with relevant guidelines and regulations and adhered to the ARRIVE guidelines. *R26^GT-Gnaq-GFP^* mice were generated using CRISPR/Cas9 gene editing using the same methodology that we previously used for generating the *R26^GT-Map2k1-GFP^* line[37], except that a p.R183Q mutant *Gnaq* cDNA was cloned into the ROSA targeting vector pR26 CAG/GFP AscI (Addgene: 74285)[38]. Founder pups were screened for incorporation of the *Gnaq* cDNA (Ex3 FP: 5’-gcacaattggttcgagaggttgatg-3’ and Ex5 RP: 5’-cattgtctgactccacaagaacttg-3’; amplicon size 405 bp). Positive pups were subsequently genotyped for 5’ and 3’ homologous recombination using primer sets with one primer located outside the arms of homology (5’ primer set: FP 5’ outside: ggctaggtaggggatcgggactctg; RP-SA: tggctggcaactagaaggcacactc. 3’ primer set: FP-GFP: tctcggcatggacgagctgtacaag; RP 3’ outside: gttctgagaccattctcagtggctc). Correctly targeted founders were mated with C57BL6/J mice (Jackson Laboratory: 000664) to establish the *R26^GT-Gnaq-GFP^* mouse line. Mice of this line are maintained on a C57BL6 /J background and routinely genotyped for presence of the *Gnaq* cDNA.

To obtain GNAQ p.R183Q expression specifically in ECs, *R26^GT-Gnaq-GFP/+^*females were crossed with either male *Tg(Cdh5-cre)7Mlia/J^+/-^* (Jackson Laboratory: 006137) or male *Tg(Cdh5-cre/ERT2)1Rha*^+/-^ mice[25, 39]. Tail DNA from embryos/pups was genotyped for *Gnaq* (Ex3-FP/Ex11-RP) and *Cre* (FP: 5’-caataccggagatcatgcaagctg-3’ and RP: 5’- aggcacattctagaaggtggacctg-3’; amplicon size 429 bp). The PCR cycle used for all routine genotyping was: 94°C 3’ => 36x (94°C15” => 61°C 15” => 72°C 30”) => 72°C 7’.

P1 pups received tamoxifen by dorsal subcutaneous injection of 100 µg of tamoxifen (20 µl of a 5 mg/ml tamoxifen (Sigma: T5648) dilution in sesame oil) or 5 µg of tamoxifen (20 µl of a 0.25 mg/ml tamoxifen dilution). Both male and female experimental mice were analyzed and no difference in phenotype was detected. Because of the phenotypic changes in double heterozygous mutants, blinding and masking was not possible for this study.

### Primary endothelial cell culture and droplet digital PCR (ddPCR)

We recovered lung ECs from 3-week-old *R26^GT-Map2k1-GFP/+^* and *Tg-Cdh5CreER^+/-^;R26^GT-Map2k1-GFP/+^* mice using published methods[40]. Cells were cultured for 5 days in EGM2 medium (Lonza: CC-3162) containing 1 µM of 4-OH-tamoxifen (Biogems: 6800637). DNA was extracted using the Qiagen DNeasy kit (Qiagen: 69506) and RNA was obtained using the Monarch total RNA miniprep kit (New England Biolabs: T20105).

Detection of mutant *Gnaq p.183Q* transcripts was performed by generating cDNA with the Protoscript II kit using the oligo-dT primer (New England Biolabs: E6560) and then performing ddPCR to detect wild-type and missense mutant transcripts. ddPCR primers and probes used: FP: 5’-gagtgctacgacagacgacg-3’; RP: 5’-cctttggccccctacatcg-3’. *Gnaq* wild-type probe: 5hex/cgtgcttagagttcgagtccccactac/3IABkFQ (IDT); *Gnaq* p.R183Q probe: 56-FAM/acgtgcttagagttcaagtccccactac/3IABkFQ (IDT). PCR cycle: 95°C 10’ => 40x (94°C 30” => 60°C 60” => 72°C 30”) =>98°C 10’ (ramp time: 1.2°C/second). Reactions were performed in duplicate.

### Histopathology of tissue sections

For histopathology, brain and intestines were fixed overnight with 4% paraformaldehyde and processed for paraffin embedding. 7 µm sections were cut, and standard hematoxylin/eosin staining was performed.

### Whole mount immunofluorescence staining

Intestines and dorsal ear skin were used for wholemount staining. Intestines were flushed 4 times with 1 ml PBS (2 times from each end). Intestines were then opened up. To make the intestinal wall thin enough for use with confocal microscopy the intestinal villi were gently scraped off with a closed dissection scissor. For the dorsal ear skin thin forceps were used to separate it from the underlying cartilage. Both tissues were fixed overnight in 1% paraformaldehyde at 4°C, washed with PBS (6 x 15 minutes), and blocked overnight at 4°C with PBS containing 0.3% triton-X-100 (PBS-T) and 20% Aquablock (Abcam: ab 166952 (discontinued)). Samples were incubated overnight with primary antibodies diluted in PBS-T. The following primary antibodies were used: rat anti-CD31 (1/500, BD Pharmigen, 553370); chicken anti-GFP (1/1000 Abcam, ab13970); rabbit anti-PDGFRb (1/250, Cell Signaling Technologies (CST), 3169); rabbit anti-NG2 (1/250, Sigma, AB5320); rabbit anti-alpha-SMA (1/1000, CST, 19245); Rabbit anti-HSPG2 (1/1000, Abcam, ab315029); rabbit anti-NID 2 (1/1000, Abcam, ab14513); rabbit anti-COLL IV (1/1000, Abcam, ab6586); sheep anti-ANGPT2 (1/250, R&D Systems AF7186), Goat anti-ESM1 (1/200, R&D Systems AF1999); rabbit anti-ERG (1/250, Abcam, ab92513), and rat anti-Ki67 (1/200, Invitrogen, 14-5698-82). Following the overnight incubation, samples were washed with PBS-T (3 x 40 minutes), incubated overnight at 4°C with secondary antibodies diluted in PBS-T. The following secondary antibodies were used: donkey anti-rabbit IgG-rhodamine red (1/500, Jackson ImmunoResearch, 711-295- 152); donkey anti-rat IgG-rhodamine red (1/500, Jackson ImmunoResearch, 712-295- 150); donkey anti-chicken IgG-FITC (1/500, Jackson ImmunoResearch, 703-095-155); donkey anti-rat IgG-Cy5 (1/500, Jackson ImmunoResearch, 712-175-153), donkey anti-Goat Cy5 (1/500, Jackson ImmunoResearch, 705-175-147) and donkey anti-sheep IgG-rhodamine red (1/500, Jackson ImmunoResearch, 713-295-147). Samples were then washed 1x with PBS-T containing 5 µM Hoechts 33342 (AAT-Bioquest, 17535), followed by 2 washes (40 minutes) with PBS-T). Samples were mounted with DAPI Fluoromount G (Southern Biotech, 0100-20).

For determining the EC proliferation rate, the percentage of Ki67+ ECs was calculated for ERG+/GFP- and ERG+/GFP+ ECs from 3 *Tg-Cdh5CreERT2^+/-^;R26^GT-Gnaq-GFP/+^* (DH) (using 4 to 6 single plane 20x confocal images/mouse) and 2 *R26^GT-Gnaq-GFP/+^*control animals (using 4 to 5 single plane 20x confocal images/mouse). Statistical analysis was done using the nonparametric Wilcoxon rank sum test, to account for non-normality in the groups, as assessed by the Shapiro-Wilk test. Two-tailed values of p < 0.05 were considered statistically significant. Analysis of the data was performed using Stata software version 18 (Stata Corp LLC, College Station, Texas).

### Imaging

Images of histological brain and intestine sections were taken with NIS-elements AR 3.2 software (Nikon), using a Nikon Eclipse 80i microscope equipped with a Nikon digital sight DS-Ri1 camera. Whole mount images of intestine, brain and fluorescent images of the brain surface were taken with a Nikon SMZ18 stereomicroscope equipped with a Nikon digital sight DS-U3 camera using NIS-elements AR 4.4 software. Confocal images of the intestines and the dorsal ear skin were taken using a Zeiss LSM 800 Confocal Microscope and Zen Blue software version 2.5. Images of control and mutant samples were adjusted identically for brightness, contrast, and sharpness (Adobe Photoshop 2025).

## ACKNOWLEDGMENTS

We thank the IDDRC Gene Manipulation & Genome Editing Core for generation of the ROSA-GT-*Gnaq*-R183Q mouse line. We also thank Dr. William T Pu, Department of Cardiology, Boston Children’s Hospital for providing the *Tg(Cdh5-cre/ERT2)1Rha+/-* mouse line.

## STATEMENTS AND DECLARATIONS

Research reported in this publication was supported by: NIH (NHLBI)-2R01HL127030-05 (AKG, JB), NIH (NICHD)-T32 HD104582-01A1 (MA), Taylor/McDonald Family, and P50 HD105351 (IDDRC Gene Manipulation & Genome Editing Core Boston Children’s Hospital). The content of this research is solely the responsibility of the authors and does not necessarily represent the official views of the National Institutes of Health.

## DATA AVAILABILITY

The *R26^GT-Gnaq-GFP^*mouse line generated for this study will be available to other researchers upon request and will be donated to Jackson Laboratory.

## REFERENCES

1. Jacobs, A.H., and Walton, R.G. (1976). The incidence of birthmarks in the neonate. Pediatrics 58, 218–222.

2. Gupta, A., and Kozakewich, H. (2011). Histopathology of vascular anomalies. Clin Plast Surg 38, 31–44.

3. Tan, W., Wang, J., Zhou, F., Gao, L., Yin, R., Liu, H., Sukanthanag, A., Wang, G., Mihm, M.C., Jr., Chen, D.B., et al. (2017). Coexistence of Eph receptor B1 and ephrin B2 in port-wine stain endothelial progenitor cells contributes to clinicopathological vasculature dilatation. Br J Dermatol 177, 1601–1611.

4. Greene, A.K., Taber, S.F., Ball, K.L., Padwa, B.L., and Mulliken, J.B. (2009). Sturge-Weber syndrome: soft-tissue and skeletal overgrowth. J Craniofac Surg 20 *Suppl 1*, 617–621.

5. Desai, S., and Glasier, C. (2017). Sturge-Weber Syndrome. N Engl J Med 377, e11.

6. Dutkiewicz, A.S., Ezzedine, K., Mazereeuw-Hautier, J., Lacour, J.P., Barbarot, S., Vabres, P., Miquel, J., Balguerie, X., Martin, L., Boralevi, F., et al. (2015). A prospective study of risk for Sturge-Weber syndrome in children with upper facial port-wine stain. J Am Acad Dermatol 72, 473–480.

7. Huikeshoven, M., Koster, P.H., de Borgie, C.A., Beek, J.F., van Gemert, M.J., and van der Horst, C.M. (2007). Redarkening of port-wine stains 10 years after pulsed-dye-laser treatment. N Engl J Med 356, 1235–1240.

8. Hammill, A.M., and Boscolo, E. (2024). Capillary malformations. J Clin Invest 134.

9. Shirley, M.D., Tang, H., Gallione, C.J., Baugher, J.D., Frelin, L.P., Cohen, B., North, P.E., Marchuk, D.A., Comi, A.M., and Pevsner, J. (2013). Sturge-Weber syndrome and port-wine stains caused by somatic mutation in GNAQ. N Engl J Med 368, 1971–1979.

10. Couto, J.A., Huang, L., Vivero, M.P., Kamitaki, N., Maclellan, R.A., Mulliken, J.B., Bischoff, J., Warman, M.L., and Greene, A.K. (2016). Endothelial Cells from Capillary Malformations Are Enriched for Somatic GNAQ Mutations. Plast Reconstr Surg 137, 77e–82e.

11. Couto, J.A., Ayturk, U.M., Konczyk, D.J., Goss, J.A., Huang, A.Y., Hann, S., Reeve, J.L., Liang, M.G., Bischoff, J., Warman, M.L., et al. (2017). A somatic GNA11 mutation is associated with extremity capillary malformation and overgrowth. Angiogenesis 20, 303–306.

12. Dompmartin, A., van der Vleuten, C.J.M., Dekeuleneer, V., Duprez, T., Revencu, N., Desir, J., Te Loo, D., Flucke, U., Eijkelenboom, A., Schultze Kool, L., et al. (2022). GNA11-mutated Sturge-Weber syndrome has distinct neurological and dermatological features. Eur J Neurol 29, 3061–3070.

13. Huang, L., Couto, J.A., Pinto, A., Alexandrescu, S., Madsen, J.R., Greene, A.K., Sahin, M., and Bischoff, J. (2017). Somatic GNAQ Mutation is Enriched in Brain Endothelial Cells in Sturge-Weber Syndrome. Pediatr Neurol 67, 59–63.

14. Bichsel, C.A., Goss, J., Alomari, M., Alexandrescu, S., Robb, R., Smith, L.E., Hochman, M., Greene, A., and Bischoff, J. (2019). Association of Somatic GNAQ Mutation With Capillary Malformations in a Case of Choroidal Hemangioma. JAMA Ophthalmol 137, 91–95.

15. Goddard, L.M., and Iruela-Arispe, M.L. (2013). Cellular and molecular regulation of vascular permeability. Thromb Haemost 109, 407–415.

16. Sandoval, Y.H., Atef, M.E., Levesque, L.O., Li, Y., and Anand-Srivastava, M.B. (2014). Endothelin-1 signaling in vascular physiology and pathophysiology. Curr Vasc Pharmacol 12, 202–214.

17. Barki-Harrington, L., Luttrell, L.M., and Rockman, H.A. (2003). Dual inhibition of beta-adrenergic and angiotensin II receptors by a single antagonist: a functional role for receptor-receptor interaction in vivo. Circulation 108, 1611–1618.

18. Martins, L., Giovani, P.A., Reboucas, P.D., Brasil, D.M., Haiter Neto, F., Coletta, R.D., Machado, R.A., Puppin-Rontani, R.M., Nociti, F.H., Jr., and Kantovitz, K.R. (2017). Computational analysis for GNAQ mutations: New insights on the molecular etiology of Sturge-Weber syndrome. J Mol Graph Model 76, 429–440.

19. Litosch, I. (2016). Decoding Galphaq signaling. Life Sci 152, 99–106.

20. Wellman, R.J., Cho, S.B., Singh, P., Tune, M., Pardo, C.A., Comi, A.M., and Workgroup, B.S.-W.s.P. (2019). Galphaq and hyper-phosphorylated ERK expression in Sturge-Weber syndrome leptomeningeal blood vessel endothelial cells. Vasc Med 24, 72–75.

21. Huang, L., Bichsel, C., Norris, A.L., Thorpe, J., Pevsner, J., Alexandrescu, S., Pinto, A., Zurakowski, D., Kleiman, R.J., Sahin, M., et al. (2022). Endothelial GNAQ p.R183Q Increases ANGPT2 (Angiopoietin-2) and Drives Formation of Enlarged Blood Vessels. Arterioscler Thromb Vasc Biol 42, e27–e43.

22. Schrenk, S., Bischoff, L.J., Goines, J., Cai, Y., Vemaraju, S., Odaka, Y., Good, S.R., Palumbo, J.S., Szabo, S., Reynaud, D., et al. (2023). MEK inhibition reduced vascular tumor growth and coagulopathy in a mouse model with hyperactive GNAQ. Nat Commun 14, 1929.

23. Wetzel-Strong, S.E., Galeffi, F., Benavides, C., Patrucco, M., Bullock, J.L., Gallione, C.J., Lee, H.K., and Marchuk, D.A. (2023). Developmental expression of the Sturge-Weber syndrome-associated genetic mutation in Gnaq: a formal test of Happle’s paradominant inheritance hypothesis. Genetics 224.

24. Solomon, C.M., M.; Singh, P.; Nemeth, Ch.; Comi, A. (2024). R183Q GNAQ Sturge–Weber Syndrome Leptomeningeal and Cerebrovascular Developmental Mouse Model. Journal of Vascular Anomalies 5.

25. Alva, J.A., Zovein, A.C., Monvoisin, A., Murphy, T., Salazar, A., Harvey, N.L., Carmeliet, P., and Iruela-Arispe, M.L. (2006). VE-Cadherin-Cre-recombinase transgenic mouse: a tool for lineage analysis and gene deletion in endothelial cells. Dev Dyn 235, 759–767.

26. Huang, J.L., Urtatiz, O., and Van Raamsdonk, C.D. (2015). Oncogenic G Protein GNAQ Induces Uveal Melanoma and Intravasation in Mice. Cancer Res 75, 3384–3397.

27. Nasim, S., Bichsel, C., Pinto, A., Alexandrescu, S., Kozakewich, H., and Bischoff, J. (2024). Similarities and differences between brain and skin GNAQ p.R183Q driven capillary malformations. Angiogenesis 27, 931–941.

28. Witjas, F.M.R., van den Berg, B.M., van den Berg, C.W., Engelse, M.A., and Rabelink, T.J. (2019). Concise Review: The Endothelial Cell Extracellular Matrix Regulates Tissue Homeostasis and Repair. Stem Cells Transl Med 8, 375–382.

29. Rauff, A., LaBelle, S.A., Strobel, H.A., Hoying, J.B., and Weiss, J.A. (2019). Imaging the Dynamic Interaction Between Sprouting Microvessels and the Extracellular Matrix. Front Physiol 10, 1011.

30. Payne, L.B., Zhao, H., James, C.C., Darden, J., McGuire, D., Taylor, S., Smyth, J.W., and Chappell, J.C. (2019). The pericyte microenvironment during vascular development. Microcirculation 26, e12554.

31. Nasim, S., Bichsel, C., Dayneka, S., Mannix, R., Holm, A., Vivero, M., Alexandrescu, S., Pinto, A., Greene, A.K., Ingber, D.E., et al. (2024). MRC1 and LYVE1 expressing macrophages in vascular beds of GNAQ p.R183Q driven capillary malformations in Sturge Weber syndrome. Acta Neuropathol Commun 12, 47.

32. Sundaram, S.K., Michelhaugh, S.K., Klinger, N.V., Kupsky, W.J., Sood, S., Chugani, H.T., Mittal, S., and Juhasz, C. (2017). GNAQ Mutation in the Venous Vascular Malformation and Underlying Brain Tissue in Sturge-Weber Syndrome. Neuropediatrics 48, 385–389.

33. Van Raamsdonk, C.D., Bezrookove, V., Green, G., Bauer, J., Gaugler, L., O’Brien, J.M., Simpson, E.M., Barsh, G.S., and Bastian, B.C. (2009). Frequent somatic mutations of GNAQ in uveal melanoma and blue naevi. Nature 457, 599–602.

34. Kimple, A.J., Bosch, D.E., Giguere, P.M., and Siderovski, D.P. (2011). Regulators of G-protein signaling and their Galpha substrates: promises and challenges in their use as drug discovery targets. Pharmacol Rev 63, 728–749.

35. Galeffi, F., Snellings, D.A., Wetzel-Strong, S.E., Kastelic, N., Bullock, J., Gallione, C.J., North, P.E., and Marchuk, D.A. (2022). A novel somatic mutation in GNAQ in a capillary malformation provides insight into molecular pathogenesis. Angiogenesis 25, 493–502.

36. Van Raamsdonk, C.D., Griewank, K.G., Crosby, M.B., Garrido, M.C., Vemula, S., Wiesner, T., Obenauf, A.C., Wackernagel, W., Green, G., Bouvier, N., et al. (2010). Mutations in GNA11 in uveal melanoma. N Engl J Med 363, 2191–2199.

37. Smits, P.J., Sudduth, C.L., Konczyk, D.J., Cheng, Y.S., Vivero, M.P., Kozakewich, H.P.W., Warman, M.L., and Greene, A.K. (2023). Endothelial cell expression of mutant Map2k1 causes vascular malformations in mice. Angiogenesis 26, 97–105.

38. Chu, V.T., Weber, T., Graf, R., Sommermann, T., Petsch, K., Sack, U., Volchkov, P., Rajewsky, K., and Kuhn, R. (2016). Efficient generation of Rosa26 knock-in mice using CRISPR/Cas9 in C57BL/6 zygotes. BMC Biotechnol 16, 4.

39. Wang, Y., Nakayama, M., Pitulescu, M.E., Schmidt, T.S., Bochenek, M.L., Sakakibara, A., Adams, S., Davy, A., Deutsch, U., Luthi, U., et al. (2010). Ephrin-B2 controls VEGF-induced angiogenesis and lymphangiogenesis. Nature 465, 483–486.

40. Wang, J., Niu, N., Xu, S., and Jin, Z.G. (2019). A simple protocol for isolating mouse lung endothelial cells. Sci Rep 9, 1458.

